# Exploring the microbiome of black soil from Central India and the impacts of agricultural practices on soil microbial communities

**DOI:** 10.1101/2024.12.29.630697

**Authors:** Sudhir Kumar, Rituja Saxena, Rajesh Saxena, Vineet K Sharma

## Abstract

**Background:** Healthy microbial communities in soil are indispensable for agricultural productivity. The diversity in physiochemical properties, their geological origins, and local climatic conditions lead to the evolution of diverse soil types with distinct bacterial communities. Despite a global distribution, the microbial composition of black soil remains largely unexplored. Here, we investigate the composition, diversity, and functional potential of bacterial communities in black soils from central India. We also examine how common agricultural disturbances like fertilizer application and the growth of crops impact soil communities in a three-year longitudinal field experiment involving three crops.

**Results:** The black soil core microbiome was dominated by phyla Actinobacteriota and Proteobacteria members. Analysis of bacterial composition in vermicompost revealed the presence of plant growth-promoting taxa. Among the assessed cropping practices, crop growth significantly affected microbiome composition in soil. We also observed the impact of crops in selectively enhancing the microbiome in the soil, highlighting the role of plant species in shaping community structure. During this experiment, fertilizer treatment did not influence the microbiome significantly. Overall, the data showed the vulnerability and resilience of soil bacterial communities to agricultural disturbances.

**Conclusion:** The microbial composition of black soil in central India was revealed, and the effects of agricultural management on black soil microbial composition were evaluated. The prominent impact of individual plants showed the role of plant-root microbiome interactions in shaping the composition of soil communities. Fertilizer application did not show significant effects during the field experiment, reflecting the resilience of soil communities to short-term disturbances. We also shed light on the functional metagenomic potential of black soil communities and the effect of common agricultural disturbances in a prominent soil type.

## Background

The enormous diversity of soil bacterial communities is critical to environmental health. It influences agricultural productivity and even the response to climate change. It can also impact human health by serving as a reservoirs of antimicrobial resistance and pathogens (Whitman et al. 1998; Banerjee and van der Heijden 2023; Yang et al. 2024). This diversity of microbial communities results from immense spatial and temporal variability of physicochemical properties in soils (Philippot et al. 2023). However, previous work has identified pH, organic carbon, oxygen, moisture, nitrogen, and phosphorus as factors that influence the microbial composition of soils (Fierer 2017). The considerable geographical variation in soils requires detailed information from diverse regions and soil types worldwide.

The soil type, a combined effect of the parent material, climatic conditions, living organisms, and physicochemical properties, has recently become important in understanding variation in soil microbiome. Studies have explored other regional environments, but various Indian soil types remain understudied (Mittal et al. 2019; Prasoodanan P. K et al. 2024).

Black soil, also known as Regur or black cotton soil, is distributed globally across Asia, Africa, Australia, and Europe, but its microbial composition remains unstudied (S R and K Y 2013; Pershina et al. 2018). The climate of the sampled area is sub-humid and subtropical, with hot summers and mild winters. The soil originates from the parent material of basalt rocks, has high clay content and organic matter, and is classified as typic Haplustert of Vertisols (Mandal et al. 2013; Mohanty et al. 2014). These factors make Indian black soil distinct in its geological origin and biological properties from other counterparts. In India, it is a significantly important group among 11 largely unexplored soil types in the subcontinent. Distributed in the states of Maharashtra, Madhya Pradesh, Gujarat, and Andhra Pradesh, it represents more than 70 million hectares, roughly constituting 21% of the total geographical area of India (Chandran et al. 2012; Bhattacharyya et al. 2013). Black soils are deep to shallow, dark-colored soils with distinct clay mineralogy and a high amount of organic carbon, calcium carbonate, and magnesium but are deficient in nitrogen and phosphorus. They are often difficult to cultivate due to poor subsoil porosity and aeration. Black soil is suitable for cultivating cotton, wheat, soybean, pigeon pea, groundnut, and chickpeas, which are important crops in the sampled area (Pal et al. 2012; Velmourougane et al. 2014). Due to their expanse and economic importance in Indian and global agriculture, there is a need for a comprehensive examination of microbial communities in black soil (Velmourougane et al. 2014; Pershina et al. 2018). Moreover, with the increasing interest in soil health, microbial diversity, and agricultural productivity, it is imperative to study the dynamics of microbial communities in different soil types (Hartman et al. 2018; Kumar et al. 2018; Wang et al. 2020). Such studies can also identify sensitive and rapid markers of land use-related effects on soil.

Previous work has explored the impacts of management practices like tillage, fertilizer application, pesticides, and organic farming on the microbiome in short and long-term field experiments (Hu et al. 2011; Williams et al. 2013; Balota et al. 2014; Cassman et al. 2016; Hartman et al. 2018; Gupta et al. 2022). Organic farming is considered a sustainable alternative to fertilizer use. However, organic fertilizers can also influence and shift taxonomic composition and functional potential in soil microbiome (Reganold and Wachter 2016; Kumar et al. 2018). The changes due to the crop species under cultivation also need to be evaluated to understand the effect of monoculture farming on soils. Multiple studies have explored the microbial diversity in Indian soils and different agricultural practices, but none have focused on black soil (Rao et al. 2016; Kumar et al. 2019; Chaudhari et al. 2020; Suyal et al. 2021).

To address this knowledge gap, we conducted a longitudinal field experiment and utilized bacterial 16S amplicon sequencing to study the bacterial community structure of black soil in central India. During this field experiment, we examined the changes in soil microbial diversity and compositional shifts due to organic and chemical fertilizer-based treatments and growth for three different plant species for three years.

## Materials and Methods

### Field experiment design and soil collection

The experiment was conducted in an agricultural field (150X60 ft) in central India. The soil was high clay content, typic Haplustert of Vertisols, widespread in central India. The plot was divided into nine blocks (30X15 ft) with appropriate separation to avoid cross-contamination. The field conditions were similar for all plots before starting the field trial. Crops grown in the same season were chosen to prevent the effects of seasonal variation on soil microbiome. We cultivated the wheat variety Lok-1 (*Triticum aestivum*), linseed or flax seed local variety (*Linum usitatissimum*), and chickpea variety JG-322 (*Cicer arietinum*) for this experiment. Seeds were sown in October-November, and crops were harvested in February-March (Rabi season). All crops were subjected to the same management conditions to avoid confounders. The crops were treated with organic vermicompost, chemical fertilizer, and control (without fertilizer). The recommended dose of DAP (Diammonium Phosphate), which provided 18% N and 46 % P_2_O_5,_ was used in chemical fertilizer group subplots. Vermicompost was produced at the MPCST (Madhya Pradesh Council of Science & Technology) facility for local farm usage and was used as an organic fertilizer.

The soil samples were collected as described by Lupatini et al. (Lupatini et al. 2017). Briefly, multiple soil samples were collected from different portions of each plot and pooled into a single sample to cover the complete field, thereby covering the maximum microbial diversity represented in the field. In this manner, about 100 gm of soil was collected from each plot at 15-20 cm depth. We collected samples twice per crop, once before sowing crops and immediately before harvesting (i.e., when the crop was ready to be harvested). We followed this sampling schedule for three consecutive annual crop seasons for three years for all treatment groups and crops (**Figure 1, Supplementary Figure 1**). Vermicompost samples from batches used as fertilizer were also collected over three years. Samples were brought to the laboratory at 4°C and kept at −20°C before immediately processing them.

**Fig. 1:**
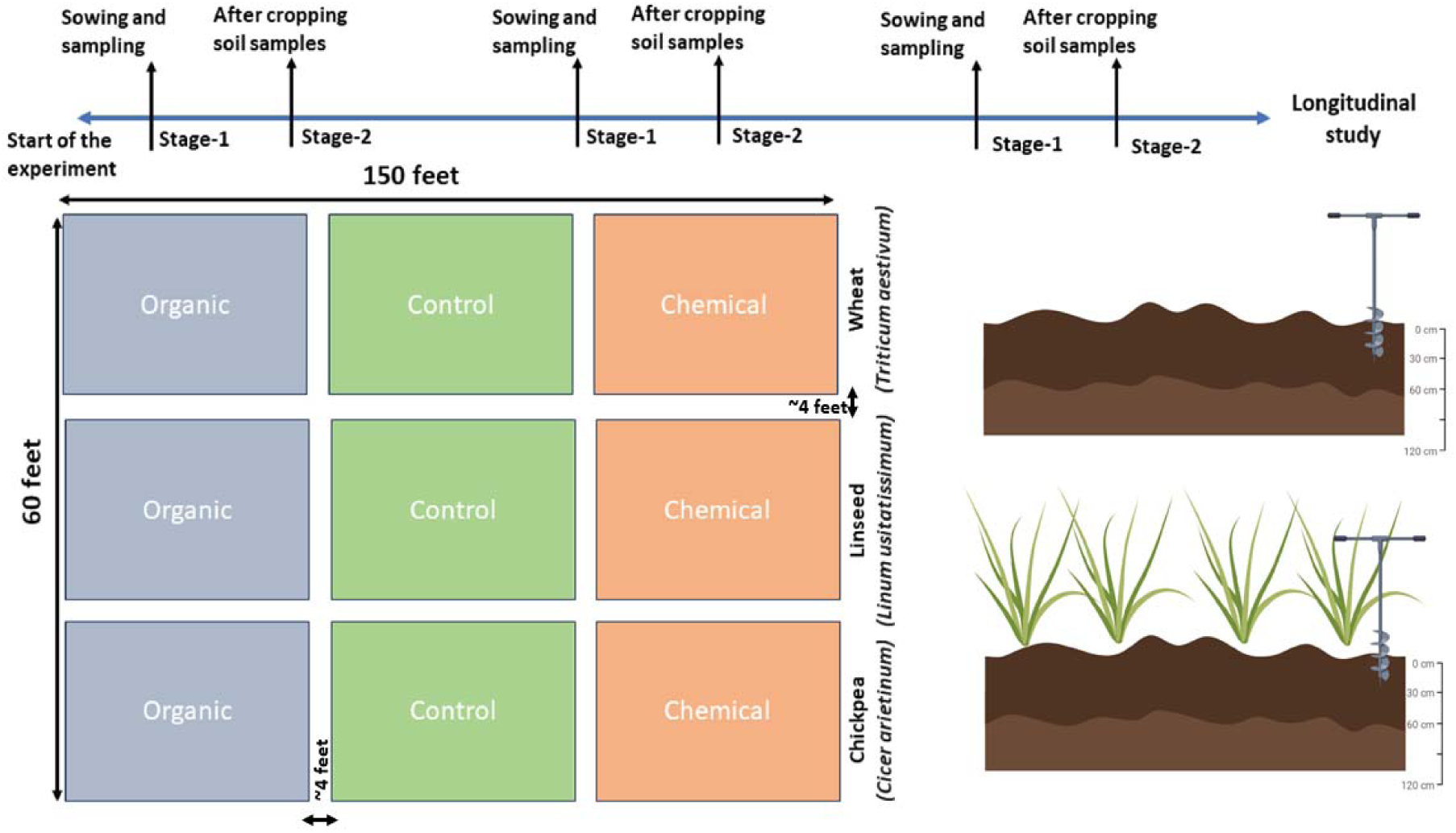
The timeline of sampling during the three years of the experiment and the design of the field experiment. Soil samples were collected for each plot before cropping and at the time of harvest.

### Soil physicochemical parameters measurement

We also measured chemical parameters in soil like the pH, organic carbon content (% oxidizable organic carbon), available phosphate as P_2_O_5_ (Kg/ha), ammonical Nitrogen as NH_3_–N (Kg/ha), available potassium (K_2_O), length of root and shoot for crops (cm), and yield (Kg/ha) for each soil sample using Soil Testing Kit for Macronutrients (Himedia Laboratories, Mumbai, India).

### DNA extraction, library preparation, and sequencing

Metagenomic DNA isolation was carried out for all soil samples using Fast DNA Spin Kit for Soil (MP Biomedicals LLC, CA, USA) using the manufacturer protocol with minor modifications. The V3 region of the bacterial 16S rRNA gene was amplified using the 341F-534R primer pair from each soil metagenomic DNA sample, and next-generation sequencing on the Illumina platform was performed after library preparation (Saxena et al. 2017).

### Read processing and data analysis

The quality filtration was done using the NGSQC toolkit, and no ambiguous bases were allowed. Reads with 80% bases below Q25 were filtered out (Patel and Jain 2012). Primer sequences were trimmed using cutadapt (Martin 2011), and high-quality paired-end reads were imported into QIIME2 for further processing. DADA2 was used for denoising and chimera filtering, and the resulting ASV (Amplicon Sequence Variants) table was filtered to remove low-abundant features and used for all downstream processing. (Callahan et al. 2016; Bolyen et al. 2019). Alpha and beta diversity analysis was carried out in QIIME 2. To account for the variation in sampling depth, we randomly subsampled the reads without replacement to the depth of the smallest sample in the dataset. Taxonomic assignment of ASVs was done using the q2-greengenes2 plugin with the latest Greengenes2 database and normalized to percent relative abundance (McDonald et al. 2023). PICRUSt-2, a tool for predictive functional profiling of amplicon data, was used to predict the functional potential of communities (Douglas et al. 2020). To determine the core microbial composition of each sample group (treatment and crops), we selected genera that were present in all samples in the group and had a minimum relative abundance of >0.001%.

### Statistical analysis

Differentially abundant taxa in groups were identified using LEfSe, which combines linear discriminant analysis and the Wilcoxon or Kruskal-Wallis test (Segata et al. 2011). Statistical comparisons and plotting were carried out in R. PERMANOVA was implemented in R using the adonis2 function with 999 permutations. Venn diagrams were constructed using web tool available at https://bioinformatics.psb.ugent.be/webtools/Venn/.

## Results

### Sample collection and sequence data processing

During three years of field experiment, 49 samples from three crops and three fertilizer treatment groups were collected to assess the bacterial community composition, including three vermicompost samples (**Figure 1)**. Sequencing of the V3 region of the 16S rRNA gene resulted in 33,868,962 (705,603.37 ± 400,358.95, mean ± sd) paired-end reads from 48 samples to analyze community composition. At this stage, one sample was excluded due to low read count. After the quality filtration of the sequence data, 25,098,680 (522,889.16 ± 293,474.85) high-quality reads remained, which were used for further processing.

Paired-end reads (25,098,680) were imported into QIIME 2 for further processing. Denoising and chimera filtering in DADA2 resulted in 30,209 ASVs. Additional filtering of low-abundant ASVs resulted in 24,237 ASVs with a cumulative frequency of 20,822,744. Rarefaction analysis revealed sufficient sequencing depth for all samples.

### Composition and diversity of core microbiome of black soil from central India

We selected a representative set of black soil samples after excluding samples collected after cropping to avoid the effects of crop growth. The samples that received chemical fertilizer were also excluded. The black soil core microbiome represented bacteria from 14 phyla with variable relative abundances. Actinobacteriota and Proteobacteria (42.72% and 24.49%, respectively) dominated the core microbiome. These were followed by Acidobacteriota, Firmicutes_D, and Chloroflexota. At the family level, there were 67 families in the core microbiome. These include Rubrobacteraceae, Geodermatophilaceae, Micromonosporaceae, Pyrinomonadaceae_433871, and JACDCH01 (order Acidimicrobiales) as the most highly abundant groups.

Black soil core microbiome consisted of 92 genera, unidentified *Rubrobacteraceae* (*f Rubrobacteraceae;g*) was the most abundant group, followed by *Geodermatophilus_A*, *PSRF01* (Pyrinomonadaceae_433871), *VFJN01* (order Acidimicrobiales family JACDCH01), and *Palsa-739* (family Gaiellaceae).

### Alpha diversity in black soil microbiome

Alpha diversity in black soil was quantified using Faith’s PD, Shannon entropy, Pielou’s evenness, and the number of observed features (ASVs). The average Shannon entropy was 9.80 ± 0.32 (average ± sd), and there were 2,777.5 ± 670.69 (average ± sd) ASVs. These results indicate a significant degree of variety and evenness, representing variation in diversity and composition among the chosen black soil samples. (**Figure 2)**.

**Fig. 2:**
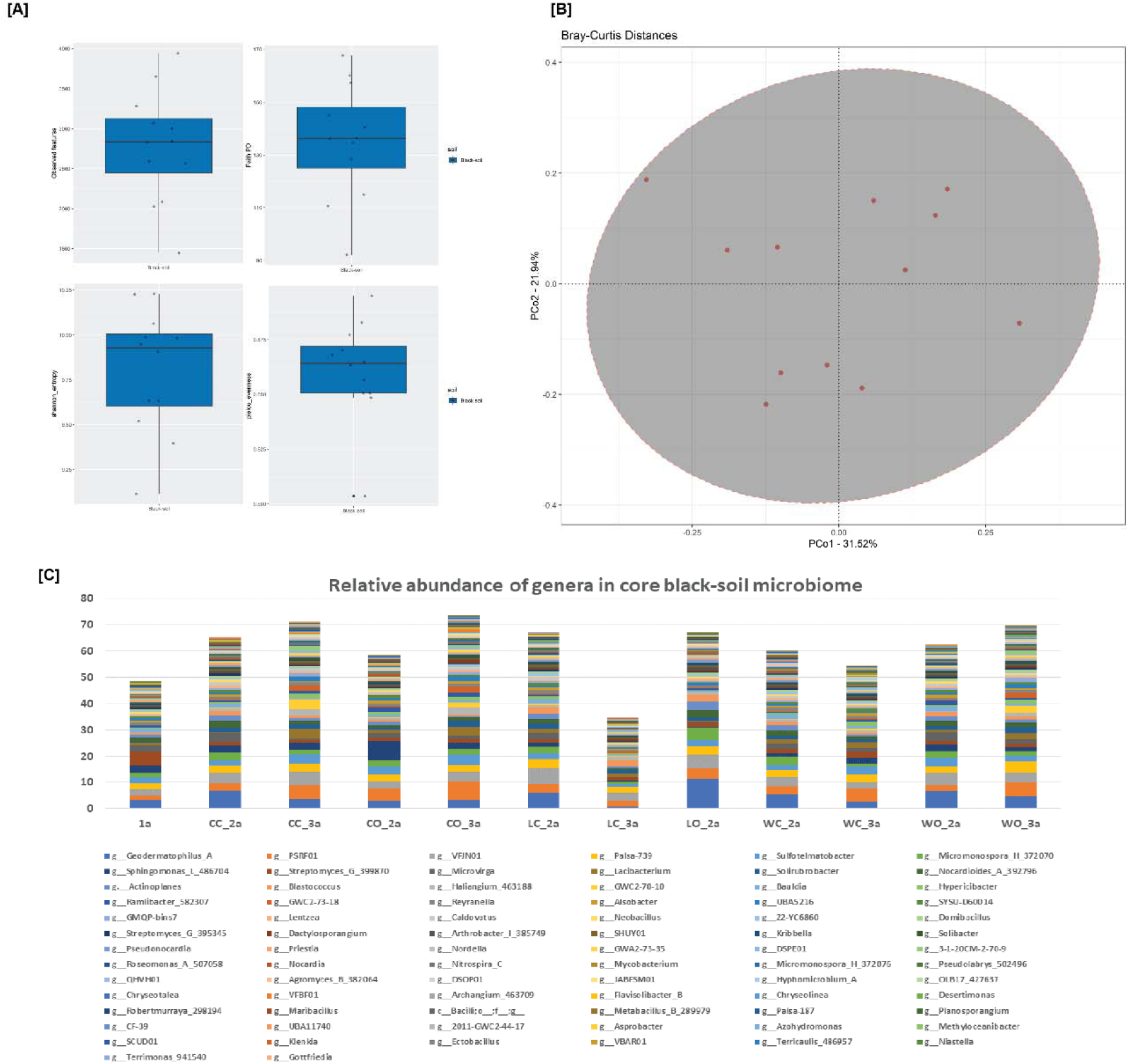
**[A]** Box plots representing alpha diversity in black soil samples. **[B]** Principal coordinate plot of Bray-cutis dissimilarity between black soil samples showing variation in black soil composition. **[C]** Relative abundance (%) of genera in the core microbiome of black soil samples.

### Taxonomic composition of vermicompost samples

PCoA revealed a distinct clustering of vermicompost samples away from soil samples, suggesting a different composition. In vermicompost samples, phylum Actinobacteriota was the most abundant, followed by Proteobacteria, Firmicutes_D, Firmicutes_A, and Bacteroidota. At the genus level, the most abundant group was *Bacillus_H* (Bacillaceae_D_361233,), followed by *Paraclostridium*, *Streptomyces_G_395345*, *Kribbella*, and *Nocardioides_A_392796*.

### Effect of cropping on the diversity of soil microbiome

A comparison of microbial composition between before and after cropping samples revealed the impact of cropping on the soil microbiome. The phylogenetic diversity (Faith’s PD) was significantly higher in the after-cropping group. A similar (non-significant) trend was observed for the number of observed features and Shannon entropy (**Figure 3-C**).

**Fig. 3:**
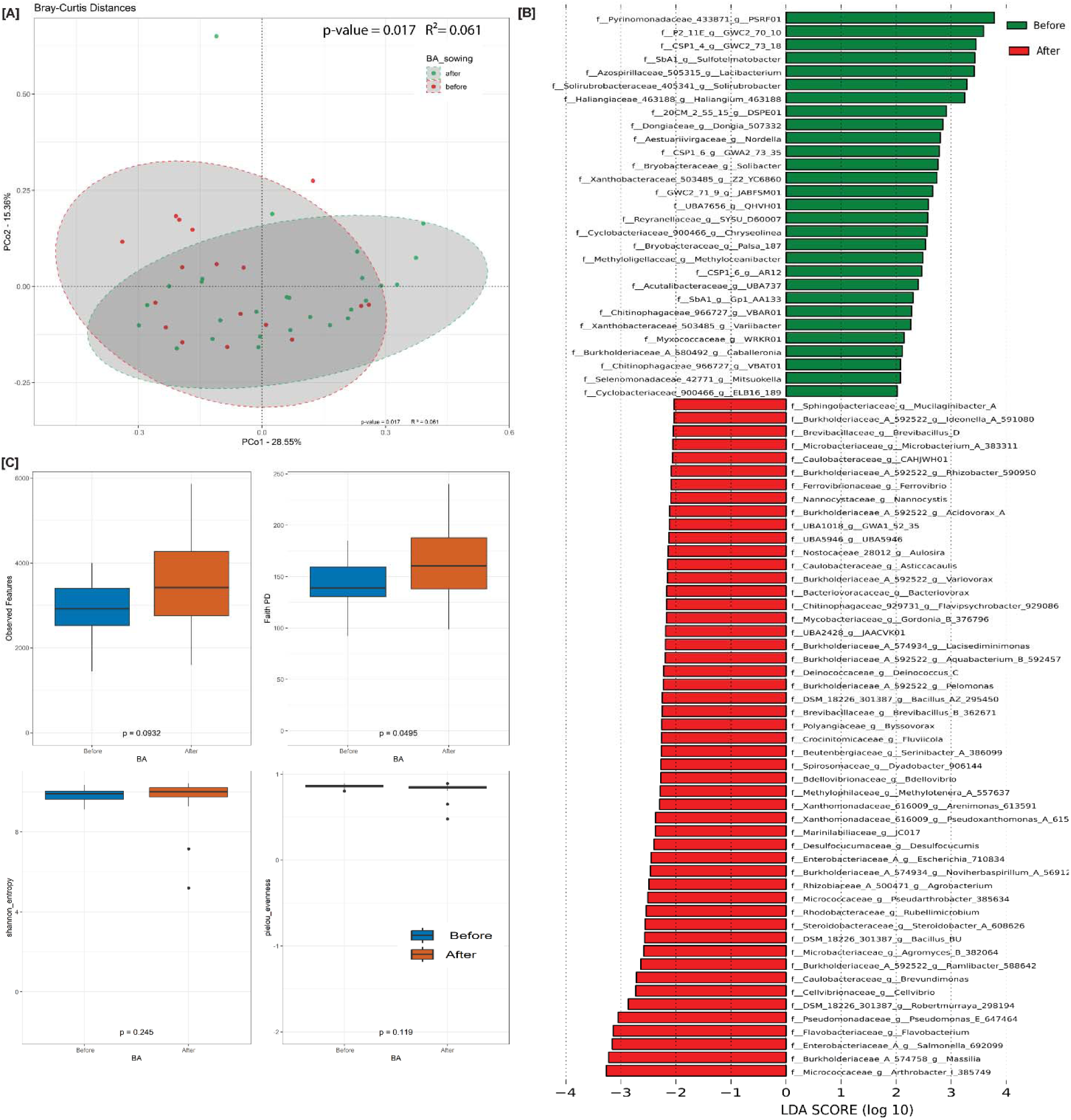
**[A]** Principal coordinate plot based on Bray-Curtis dissimilarity between soil samples under cultivation, samples are colored based on before and after cropping groups. **[B]** Differentially abundant genera in before and after-cropping groups. **[C]** Alpha diversity in before and after-cropping groups.

The alpha diversity values in fertilizer treatment groups did not differ, suggesting no significant effect on microbial diversity between these groups. There was no significant correlation between the soil chemical properties and alpha diversity.

Differences in the community composition between samples (beta-diversity) were analyzed using Bray-Curtis inter-sample dissimilarity plotted as principal coordinate plots. A significant difference was observed between before and after-cropping soil samples irrespective of the type of crop or fertilization treatment they were subjected to, indicating changes in microbiome composition as a result of cropping. The first two principal coordinates explained 28.55% and 15.36% variation (PERMANOVA p= 0.017, R^2^ = 0.06) (**Figure 3-A)**. There was a significant difference in the type of crop under cultivation (PERMANOVA p=0.027, R^2^ =0.08), showing the effect of individual plant species on the microbial composition of the soil (**Figure 4-C)**. There was no significant effect of fertilizer treatment groups on beta diversity (**Figure 5-C)**.

**Fig. 4.**
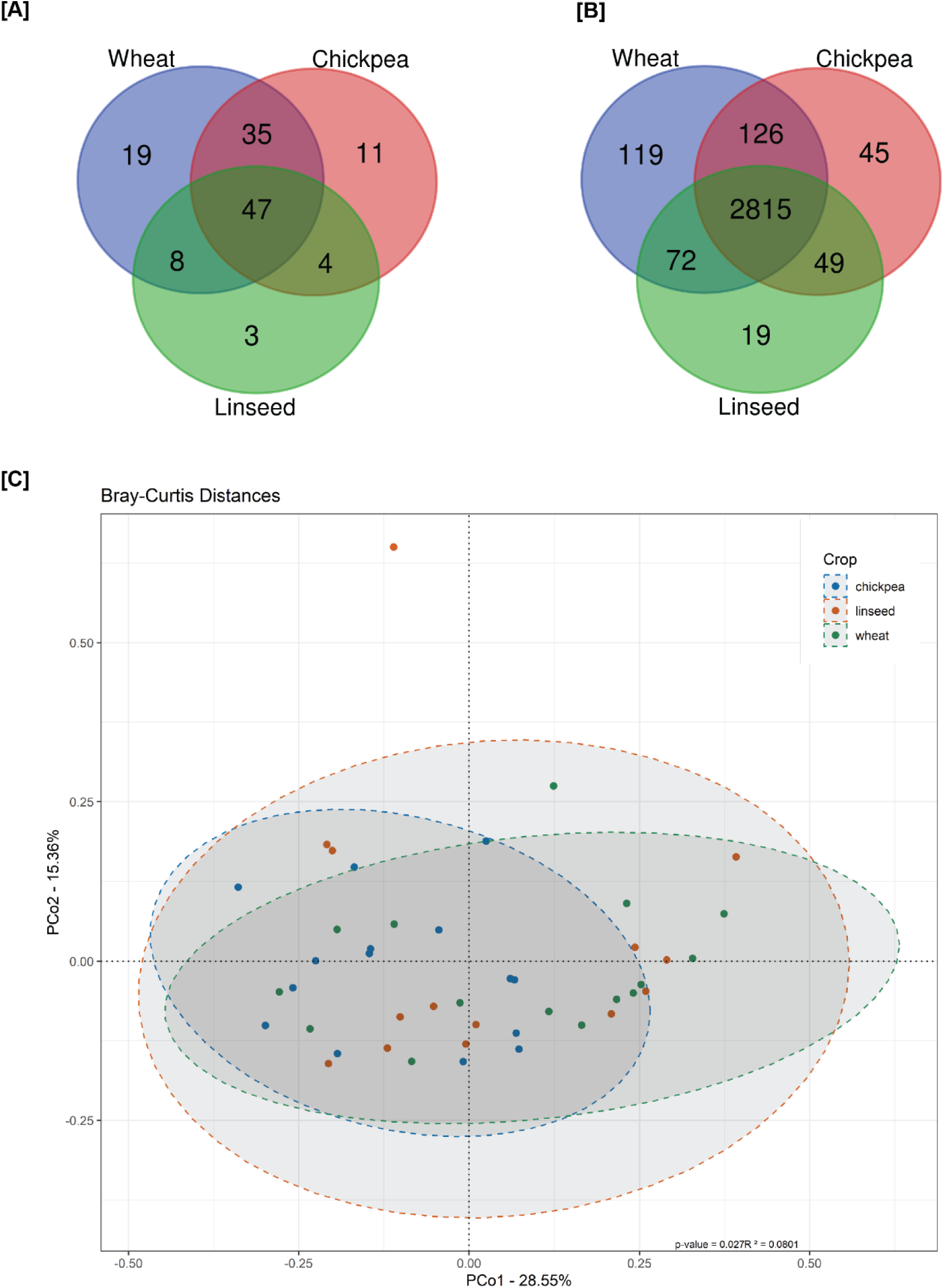
**[A]** Venn diagram showing unique and shared genera in cropping groups. **[B]** Venn diagram showing unique and shared KOs in cropping groups. **[C]** Principal coordinate plot based on Bray-Curtis dissimilarity between soil samples grouped by crop under cultivation.

**Fig. 5.**
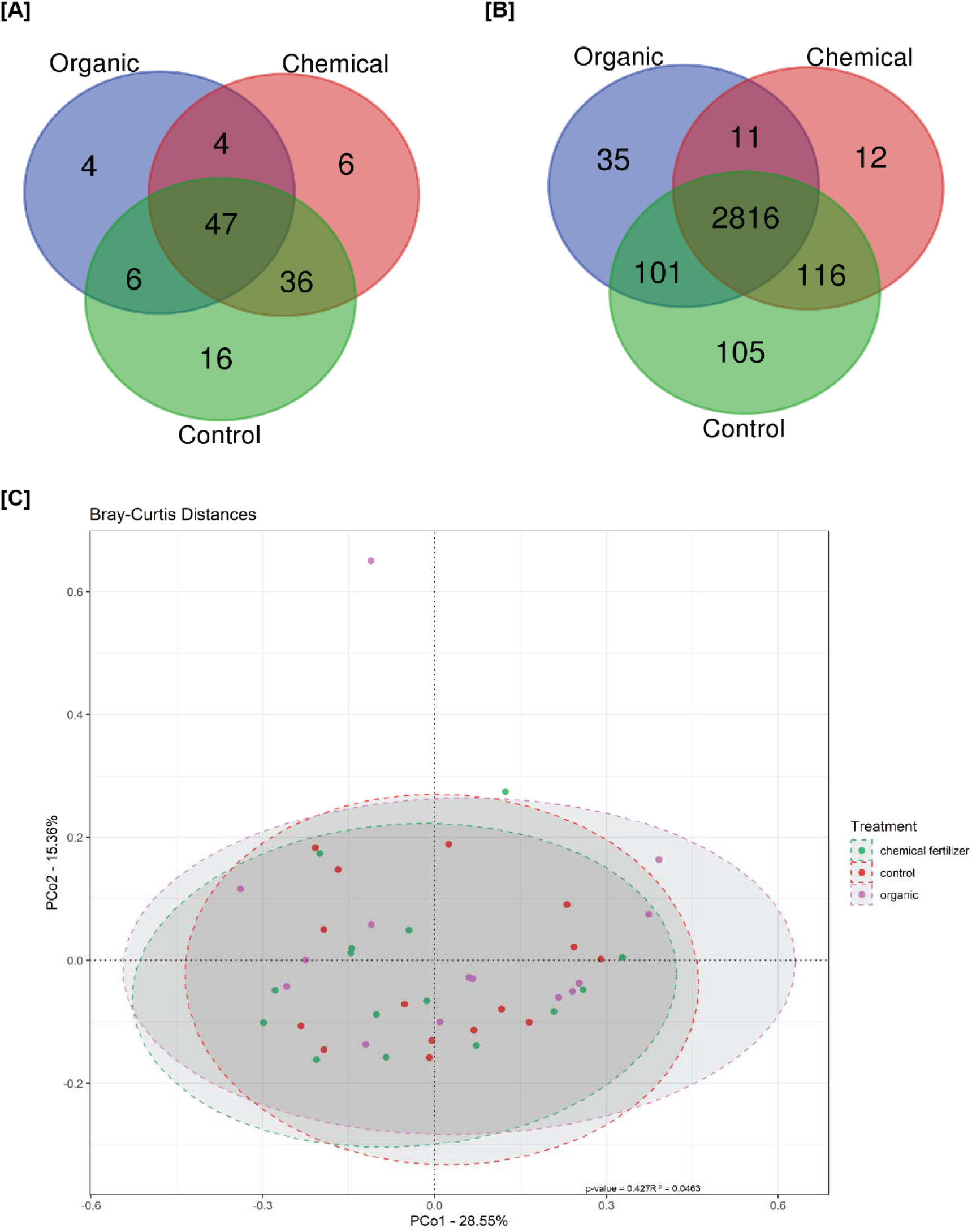
**[A]** Venn diagram showing unique and shared genera in fertilizer treatment groups. **[B]** Venn diagram showing unique and shared KOs in fertilizer treatment groups. **[C]** Principal coordinate plot based on Bray-Curtis dissimilarity between soil samples grouped by fertilizer treatment.

### Compositional shifts in soil microbiome due to cropping

We used LEfSe and identified genera that are differentially abundant in before and after cropping groups to identify bacterial genera affected by cropping on the land. In the before group, 31 genera from diverse bacterial families like Xanthobacteraceae_503485, Burkholderiaceae_A_580492, Chitinophagaceae_966727, and Cyclobacteriaceae_900466 etc., were at higher abundance. Whereas, in the after-cropping group, Burkholderiaceae, Caulobacteraceae, Xanthomonadaceae_616009, Brevibacillaceae, Microbacteriaceae, and Nostocaceae, among others were identified (**Figure 3-C).**

### Changes in soil microbiome composition with crop species and fertilizer groups

Venn diagrams were constructed to identify shared and unique genera in the defined groups. We found 109, 62, and 97 in the wheat, linseed, and chickpea-cultivated soil group. All groups shared 47 genera common to all. However, the wheat, linseed, and chickpea-cultivated groups had 19, 3, and 11 genera unique in their core microbiome, respectively (**Figure 4-A).**

The genera unique to wheat-cultivated soils were *Palsa-187*, *Cytobacillus_298193*, *Gottfriedia*, *Lysinibacillus_304693*, and *Vicinamibacter,* among others. Multiple Rhizobiales (*Devosia_A_502124*, *Pseudolabrys_502496*, *Afipia*, *Tepidamorphus*, and *Phreatobacter*) were unique to the chickpea group. Linseed group had *Archangium_463571*, *Sorangium*, and *Ohtaekwangia* as unique genera to the soil group.

Similarly, the control group had 105, 93, and 61 genera were found in the core microbiome of the control, chemical fertilizer, and organic fertilizer-treated group, respectively. Out of these, 16 were in the control group, six were in the chemical group, and four unique genera were in the organic fertilizer treatment group. (**Figure 5-A).**

The unique genera in the organic-treated group were *Phenylobacterium*, *Cytobacillus_298193*, *Koribacter*, and *Roseomonas_A_507058*. In the chemical fertilizer group *JABFSM01* (order Gemmatimonadales, family__GWC2-71-9), *Aridibacter*, *Tsukamurella*, *Noviherbaspirillum_A_568104*, *Tumebacillus_A*, and *Phycicoccus_A_390393* were unique. The control group included *Modestobacter*, *Gottfriedia*, *Dongia_507332*, *Sorangium*, *Pseudolabrys_502538*, *Lysinibacillus_304693*, *Skermanella*, *Pseudolabrys_502496*, *Klenkia*, and *Afipia* among others (**Figure 5).**

### Predicted functional profiles in microbial communities

The functional profiling using PICRUSt-2 revealed diverse functions in black soil samples. The most abundant functions included metabolic pathways, biosynthesis of secondary metabolites, microbial metabolism in diverse environments, biosynthesis of amino acids, quorum sensing, and carbon metabolism-related pathways. The before and after groups didn’t show any significantly different functions (KO abundance), although a number of taxa showed significant p-values in Krukal-Wallis comparisons, the effect size was not large enough to differentiate between the two groups.

MetaCyc pathway abundance data revealed a higher abundance of 21 pathways in the before-cropping group. These included diverse metabolic functions like amino acid synthesis (L-isoleucine biosynthesis IV, L-glutamate and L-glutamine biosynthesis), carbohydrate metabolism (Calvin-Benson-Bassham cycle, pentose phosphate pathway (non-oxidative branch), and GDP-D-glycero-α-D-manno-heptose biosynthesis). In comparison, the after-cropping group showed substrate degradation (superpathway of ornithine degradation, formaldehyde assimilation II (RuMP Cycle), glycine betaine degradation I), stress response regulation (ppGpp biosynthesis), adenosylcobalamin biosynthesis from cobyrinate a,c-diamide I, peptidoglycan biosynthesis IV (*Enterococcus faecium*), superpathway of thiamin diphosphate biosynthesis II, superpathway of glucose and xylose degradation, and formaldehyde oxidation I in higher abundance.

### Functions in crops and fertilizer treatment groups

We also calculated the relative abundance of KEGG orthologues in all samples and identified core functions in each group using the same criteria for genus-level analysis. In crop types, wheat showed the largest number of unique KOs followed by chickpea and linseed. The patterns observed by the shared and unique KOs were similar to genus-level comparisons, suggesting taxonomic and functional uniqueness in the soil microbiome of each group. Similarly, in the fertilizer treatment group, the control group had the most unique functions (105), followed by organic treatment (35) and the chemical treatment group (12). Similar patterns of distinctive features in the control group were observed for the predicted functional profiles (**Figure 4-B, 5-B).**

## Discussion

In this work, we have presented sequencing and analysis of bacterial 16S rRNA amplicon data from a field experiment designed to understand the composition and dynamics of black soil microbial communities and assessed the impact of agriculture management in a three-year field experiment. We evaluated the effect of chemical and organic fertilizer treatment on soil microbiomes in three different crop species for three consecutive years. The experiment was conducted in Bhopal (central India), where black soils (Vertisols), a significant soil type in India, are prominent.

One of the key outcomes was the revelation of the core microbiome of black soil. The composition was dominated by members phyla Actinobacteriota and Proteobacteria. The most abundant ASV group was assigned to the unclassified genus in the family Rubrobacteraceae. Members of Rubrobacteraceae are dominant in hyperarid and saline environments (Pershina et al. 2018). The second most abundant genus, *Geodermatophilus,* is widely present in arid soils and is resistant to desiccation, radiation, temperature changes, and ultraviolet lights (Castro et al. 2018). Rapid drying and hot climate in the sampled location can lead to conditions selecting for these genera in the soil.

Additionally, we sequenced vermicompost samples used as organic fertilizer. Vermicompost helps to improve the number of plant growth-promoting bacteria in the soil, adding to organic matter and nutrients (Pathma and Sakthivel 2012; Andleeb et al. 2022). The microbial composition of vermicompost depends on several factors, including the initial material, worm species, temperature, and moisture conditions while composting. *Bacillus*, *Streptomyces*, *Nocardioides*, and *Nitrospira* were among the most abundant in these samples and have been found in wormcast and compost-treated soil. These bacteria can potentially provide disease resistance to plants (Pathma and Sakthivel 2012; Gopalakrishnan et al. 2014; Andleeb et al. 2022). For example, *Nocardioides* can suppress the growth of *Fusarium,* which causes stem rot in tomatoes providing disease resistance (Zhao et al. 2019). Several *Streptomyces sp.* can act as plant growth promoters and protect from pathogens. In vermicompost, a higher abundance of other growth-promoting taxa, like *Paenibacillus_A*, *Pseudomonas*, *Devosia*, etc., was also noticed (Gopalakrishnan et al. 2014). In addition to the effects on nutrient availability, the seeding of beneficial bacteria from the organic fertilizers in the soil can positively impact plant growth.

The results from the comparative analysis showed that cultivation alters the microbial composition of soils and has distinct effects compared to those of aboveground plant species and fertilizer treatment. For example, a significantly higher abundance of *Massilia* in after-cropping soils can be attributed to selection due to increased cellulosic biomass in soils (Chaudhari et al. 2020). The observed increase in diversity after cropping can be attributed to plant root activity. Similar effects on soil have been reported due to continuous cropping (Wang et al. 2015; Ma et al. 2023). Root activity can affect major confounders like moisture, pH, and soil organic matter in the surrounding soil and thus is a potent modulator of soil microbiome composition (Fierer 2017). The diversity of the aboveground plant community also tends to increase overall microbial diversity in soil and is a significant determinant of soil health (Wang et al. 2020). These effects can be particularly relevant for the long-term effects of monoculture farming.

There was no significant difference in microbiome composition due to fertilizer application in treatment groups. Long-term experiments have shown a considerable effect of fertilizer treatment on soil microbiome (Williams et al. 2013; Balota et al. 2014; Kumar et al. 2018). Fertilizer application for a longer duration might exacerbate effects on the microbiome significantly. A shorter duration of the field experiment and resiliency in the soil communities can be a possible explanation. Additionally, higher organic matter, a characteristic property of black soils, can contribute to this resilience by providing a buffer for nutrition stress in communities (de Andrade Bonetti et al. 2017).

The differentially abundant taxa in the groups provide insights into specific adaptations to the type of treatment or cropping-related soil conditions. For example, several plant growth-promoting symbiotic nitrogen-fixing genera were noticed in the chickpea-grown soil. This observation also supports the effect of plant species recruiting bacteria in soil, shaping the communities in bulk soil. *Koribacter,* in chickpea-grown soil, is a nitrogen-fixing bacteria reported from plant-associated soils (Chen et al. 2019). The presence of root nodule symbionts like *Devosia_A_502124* and *Pseudolabrys_502496* in chickpea soil also supports this observation (Rivas et al. 2002; Chaudhari et al. 2020). In the linseed group, we observed uniquely present *Sorangium* (Polyangiaceae)*, Archangium_* 463571 (Myxococcaceae), and *Ohtaekwangia* (Cyclobacteriaceae) (Mohr et al. 2018; Chen et al. 2022). Myxococcota (*Sorangium* and *Archangium*) are known for their potential for biocontrol of bacterial and fungal pathogens (Chen 2024).

Comparisons of predicted functions also provided important insights into the functional composition of soil. Menaquinol synthesis, central to the respiratory process, was enriched before the cropping group. They are the most widespread respiratory quinones and have been reported to be increased upon inoculation of plant growth-promoting bacteria *Pseudomonas fluorescens LBUM677* (Jiménez et al. 2020). The pathways enriched in the after-cropping group included degradation/assimilation pathways, like ornithine degradation, formaldehyde assimilation II (RuMP Cycle), and glycine betaine degradation I, potentially indicating degradation of substrates used for nutrients and energy, as plant roots excrete diverse organic compounds into the soil. Another interesting observation in this group was the higher abundance of ppGpp biosynthesis (guanosine tetraphosphate), which plays a central role as an alarmone in signaling for adaptation to starvation and stress by modulating gene expression, nucleotide synthesis, and protein translation (Anderson et al. 2021). The undisturbed soil in the before-cropping group exhibited biosynthesis and energy generation-related functions. With the cropping disturbance, microbial community functions shifted towards substrate degradation, stress adaptation, and nutrient assimilation, possibly driven by environmental changes modulated by root activity (Berg and Smalla 2009; Tkacz et al. 2020; Byers et al. 2023).

Overall, we highlight the taxonomic and functional composition and the effects of agriculture management practices in black soils from central India. Considering that insights into functional composition come from a predictive method is essential. In the future, whole metagenome sequencing methods should be used to get detailed insights into the functional capabilities of black soil microbial communities. Following communities for more extended time periods can help reveal how long-term monoculture farming affects soil communities.

## Conclusion

This study uses a culture-independent method to present the first data on the composition and variability of microbial communities in black soil. Our results also highlight the significant impacts of agricultural practices and the role of crop species in shaping the microbiome in black soil. We also highlight the resilient nature of black soil microbial communities to short-term disturbances. In summary, the study demonstrated the sensitivity and resilience of resident bacterial communities to agricultural practices. It also enhances our understanding of microbial communities from a globally important soil type.

## Acknowledgments

SK thanks the University Grants Commission (UGC), India for the research fellowship. R.S. thanks DST-INSPIRE for research fellowship.

## Contributions

VKS conceived the work and designed the study. RTS helped in performing field trials and soil collection. RJS performed sample collection sample processing, library preparation, and sequencing work. SK, with inputs from VKS, designed the computational analysis framework and carried out data processing, statistical analysis, and interpretation of results. SK prepared the first draft of the manuscript under the supervision of VKS. All authors have read and approved the final manuscript.

## Ethics declarations

Not applicable.

